# Cerebral thickness reveals sex differences in verbal and visuospatial memory

**DOI:** 10.1101/2023.11.04.565655

**Authors:** Feng Sang, Shaokun Zhao, Zilin Li, Yiru Yang, Yaojing Chen, Zhanjun Zhang

## Abstract

Although sex differences in behavior/cognition and brain have previously been reported and reviewed, a sex difference of verbal and visuospatial memory and the underlying neural basis still need to be explored further. In this study, we used a machine learning model to explore a sex difference of the association between brain structure and verbal/visuospatial memory based on a community older cohort (n = 1153, age ranged from 50.42 to 86.67 years). We found females outperformed males in verbal memory (*t* = -5.431, *p* < 0.001), while males outperformed females in visuospatial memory (*t* = 3.201, *p* = 0.001). The key regions related to verbal memory in females include medial temporal cortex, orbitofrontal cortex, and some regions around insula. According to the overlapping with Yeo’s 7 networks, those regions are more located in limbic, dorsal attention and default model networks, and they are also associated with face recognition and perception processes. The key regions related to visuospatial memory, which is outstanding in males, include lateral prefrontal cortex, anterior cingulate gyrus, and some occipital regions. They overlapped more with dorsal attention, frontoparietal and visual networks, and associated with object recognition processes. These findings imply the memory performance advantage of females and males is related to the different memory preprocessing tendencies and its associated network.

## Introduction

Sex differences have always been a prominent topic in cognitive neuroscience (Baik et al., 2023; Lee et al., 2022; Schweitzer et al., 2023; Shanmugan et al., 2022). There are significant sex differences across many cognitive domains (Ge et al., 2021; Hirnstein et al., 2022; Lee et al., 2022), especially in verbval and visuospatial memory. Specifically, females outperfrom males in verbal-related memory while males outperfrom females in visuospatial-related memory (Brunet et al., 2020; Lee et al., 2022; Levine et al., 2016; Voyer et al., 2021). Although there are significant sex differences in brain structure and function (Ge et al., 2021; Sangha et al., 2021), the sex differences in verbal/visuospatial memory as they related to underlying brain structure are not yet well understood.

Previous studies have examined sex differences in multiple cognitive-behavioral domains (Gur et al., 2012; Herlitz & Lovén, 2013; Lippa et al., 2010; Liu et al., 2020; van de Weijer-Bergsma et al., 2022). It has been noted that females outperform males on verbal fluency (Scheuringer et al., 2017), verbal memory (Voyer et al., 2021) and episodic memory (Asperholm et al., 2019). In spatial abilities and working memory (Jonasson, 2005; Voyer et al., 2021), such as spatial navigation (especially at recalling the starting location (Persson et al., 2013)) and visuospatial memory tasks (Levine et al., 2016), males outperformed females, a feature that may be since males are better at processing and organizing information in cognitive processes, which is also reflected in tasks such as mental rotation a 3-dimensional object (Lee et al., 2022; Shepard & Metzler, 1971). However, whether this phenomenon of females outperforming in verbal abilities and males outperforming in spatial abilities exists consistently different age groups needs to be further explored.

Many studies have also reported significant sex differences in brain structure and function (Ingalhalikar et al., 2014; Sang et al., 2021; Shanmugan et al., 2022; Sowell et al., 2007). Although males brain volume is significantly larger than that of females (Liu et al., 2020), the relative gray matter volume and cerebral thickness is instead significantly larger in females than in males by normalizing with total intracranial volume (Ritchie et al., 2018; Sang et al., 2021). Moreover, the direction of sex differences differed in different regions of the brain. Previous study (Sowell et al., 2007) have reported that females have thicker gray matter cortex than males in the right inferior parietal to posterior temporal regions, and that this difference is independent of overall brain size and height. Other studies have found males show greater mean gray matter volume (GMV) of ventral occipitotemporal cortices, amygdala, putamen, and cerebellum than females, while the opposite sex difference in mean GMV is seen for superior frontal and lateral parietal cortices (Lotze et al., 2019; Ritchie et al., 2018; Ruigrok et al., 2014). However, this finding is not entirely consistent across study population characteristics, number of participants, and study design (Lee et al., 2022). Age is also an important factor affecting brain and cognition (Fjell et al., 2014; Sang et al., 2021; Zhao et al., 2019), especially in older populations. Both the brain and cognition shrink and decline to some degree with age (Fjell et al., 2014). And in some pathological changes (Y. Yang et al., 2022), this decline becomes more pronounced. In addition, education is a factor that has a significant impact on cognition and the brain (Lövdén et al., 2020; Steffener, 2021). Therefore, when considering the relationship between brain structure and cognitive abilities, the effects of other influencing factors such as age and education need to be taken into consideration.

From a methodological perspective, linear models are the traditional approach to exploring brain-behavior relationships. They are based on the presupposition that there is a linear relationship between the independent and dependent variables. However, linear models cannot account for complex interactions between brain structures and cognitions (Nelson et al., 2009; Saboo et al., 2022). Although nonlinear relationships between brain and cognition can be explored by extending the linear model, such as using the generalized linear model, this approach also still requires making assumptions about the form of nonlinear relationships prior to analysis. In addition, when using traditional analysis methods, it is necessary to consider that performing multiple statistical comparisons increases the probability of false positive results, especially since it is common to perform multiple statistical comparisons in neuroimaging analysis. In contrast, machine learning models, such as support vector machines and random forests, provide a new approach to address the problems mentioned above (Brennan et al., 2021). However, there are some problems with using these machine learning methods in cognitive neuroscience research. Among them, the accuracy and interpretability of the models are the two most important concerns for researchers (Linardatos et al., 2020; Murdoch et al., 2019; Saboo et al., 2022). An accurate model with good interpretability and without assuming the pattern of relationship between variables in advance, can help deepen our understanding of the more fundamental relationship between brain and behavior.

Based on the background discussed above, the core objective of the present study was to investigate the similarities and differences in brain structures associated with verbal memory and visuospatial memory, which are female-superior and male-superior cognitive domains respectively, between females and males in an elderly population. To this end, the present study first conducted a validation exercise to investigate the differences in two types of memory performance (verbal and visuospatial memory) and cortical thickness between females and males. Although cortical thickness and cortical area are heritable and have both distinct genetic determinants and different developmental trajectories (Ge et al., 2021), cortical thickness is more closely related to intelligence and cognitive function, and reflects the cytoarchitectural characteristics (Saboo et al., 2022). Then, a gradient boosting technique-extreme gradient boosting (XGBoost) (T. Chen & Guestrin, 2016), which was able to produce explainable and interpretable predicitons, was used to develop predictive models between cortical thickness and memory performance in male and female populations, respectively. This was done to obtain the structural brain features associated with memory performance in females and males. Finally, to summarize the dominant brain structural features corresponding to the dominant memory performance in females and males, the key brain regions related to memory performance were further localized and decoded.

## Materials and methods

### Participants

All participants in this study were the same as those in our previous study (Sang et al., 2021), were from the Beijing Aging Rejuvenation Initiative (BABRI) study (Chen et al., 2018; Yang et al., 2021), an ongoing longitudinal study examining the brain and cognitive decline in community-dwelling elderly individuals. All participants in this study met the following criteria: (1) native Chinese-speaking individuals over 50 years of age without dementia who had normal daily living abilities; (2) no history of brain tumors, neurological or psychiatric disorders, or addiction; (3) no conditions known to affect cerebral function, including alcoholism, current depression, Parkinson’s disease, or epilepsy; and (4) no contraindications to magnetic resonance imaging (MRI). Finally, 1153 participants (females/males, 738/415; age: 50.42-86.67 years) were included in the current study (as shown in Table 1). The study was approved by the Ethics Committee and Institutional Review Board of Beijing Normal University Imaging Center for Brain Research, and written informed consent was obtained from all participants.

**Table 1.** Demographic and cognitive characteristics of different sex groups.

|  | **Whole(n=1153)** | **Female(n=738)** | **Male (n=415)** |  |  |
| --- | --- | --- | --- | --- | --- |
|  | **M ± SD** | **M ± SD** | **M ± SD** | *t* | *p* |
| Demographic |  |  |  |  |  |
| Measures |  |  |  |  |  |
| Age (years) | 66.58±7.31 | 65.81±7.08 | 67.95±7.52 | -4.735 | <0.001 |
| Education (years) | 11.4±3.14 | 11.20±3.04 | 11.98±3.26 | -4.017 | <0.001 |
| Cognitive Measures |  |  |  |  |  |
| General cognition |  |  |  |  |  |
| MMSE | 27.32±2.24 | 27.41±2.21 | 27.16±2.28 | 1.783 | 0.075 |
| Memory |  |  |  |  |  |
| AVLT-total | 26.96±9.16 | 28.07±9.20 | 24.97±8.77 | 5.655 | <0.001 |
| ROCF-delay recall | 12.95±7.08 | 12.47±7.01 | 13.79±7.13 | -3.015 | 0.003 |

*Notes:* M ± SD: Mean ± Standard Deviation; MMSE, mini-mental state examination; AVLT, auditory verbal learning test; ROCF, Rey–Osterrieth complex figure test.

### Neuropsychological examination

All participants in this study were evaluated with a battery of neuropsychological tests. The Chinese version of the Mini-Mental State Examination (MMSE) served as a general cognitive function test. In this study, we focus on two kinds of memory examinations: the auditory verbal learning test (Rosenberg et al., 1984) (AVLT: total) indexing the verbal memory performance and the Rey-Osterrieth complex figure test (Tupler et al., 1995) (ROCF: delay) indexing the visualspatial memory performance.

### MRI data acquisition and preprocessing

All participants were scanned with a Siemens Trio 3.0 Tesla scanner (Siemens, Erlangen, Germany) in the Imaging Center for Brain Research at Beijing Normal University. Participants laid in a supine position with their heads fixed snugly by straps and foam pads to minimize head movement. High-resolution T_1_-weighted, sagittal 3D magnetization prepared rapid gradient echo sequences were acquired and covered the entire brain (sagittal slices = 176, repetition time (TR) = 1900 ms, echo time (TE) = 3.44 ms, slice thickness = 1 mm, flip angle = 9°, inversion time = 900 ms, field of view (FOV) = 256 × 256 mm^2^, acquisition matrix = 256 × 256).

We used the Statistical Parametric Mapping 12 (SPM12, version 7771, https://www.fil.ion.ucl.ac.uk/spm) and Computation Anatomy Toolbox 12 (CAT12, version 1932, https://neuro-jena.github.io/cat) toolbox to process the T_1_-weighted images of all participants, and reconstruction the center surface of brain and estimate the thickness of cerebral cortex. After obtaining individual thickness map, we resampled it into the common coordinate system (*fsaverage*) and smoothed with a 12 mm full width at half maximum.

Specifically, T_1_-weighted images were first preprocessed including denoising, spatial registration, bias-correction, and skull-striping. Then, the images were segmented into gray matter, white matter, and cerebrospinal fluid. After separating the whole brain into two hemispheres, removing the cerebellum and hindbrain, and filling the white matter hyper-intensities and subcortical regions, the closest voxel on the white matter boundary was estimated for every voxel in the cortical gray matter within each hemisphere as the white matter distance map. The projection-based thickness (PBT) was estimated the cortical thickness map by projecting the local maximum distance described in white matter distance map for each voxel in gray matter as the thickness of this voxel. Then, the cortical thickness map was resampled into a “*fsaverage*” template (Yotter et al., 2011) with a volume-based DARTEL (Diffeomorphic Anatomical Registration Through Exponentiated Lie Algebra) (Ashburner, 2007) to surface and smoothed with a 12 mm full width at half maximum. Finally, each hemisphere thickness was parcellated into 180 regions defined by HCP-MMP1.0 (Glasser et al., 2016) and estimate the mean thickness of each region (Figure 1a).

**Figure 1.**
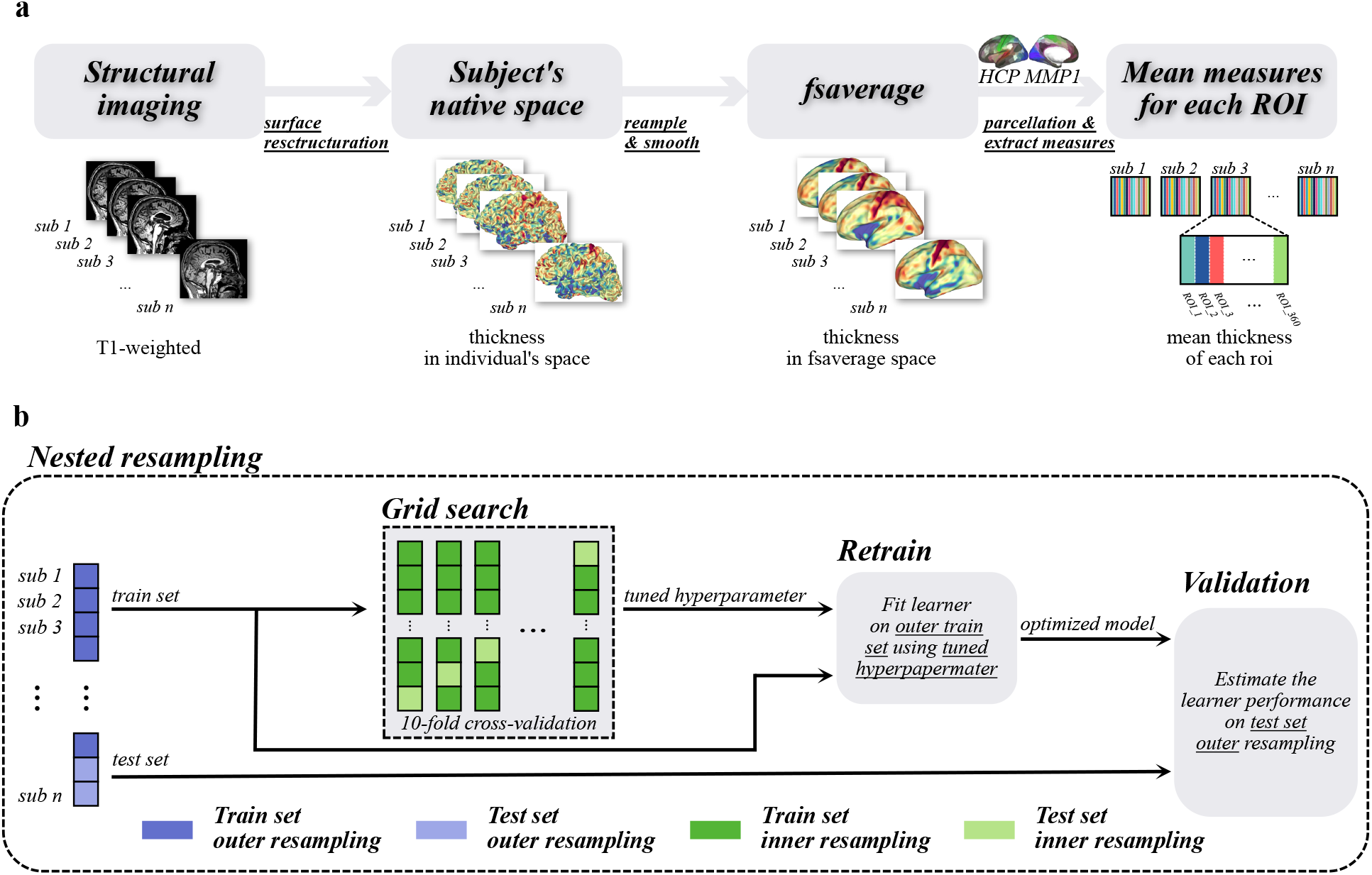
The diagram of image preprocessing and building predictive model. **a**. Each subject’s cerebral surface was reconstructed from their t1-weighted image, and to estimate the thickness in subject’s native space. As follow, the thickness map is resampled and smoothed into *fsaverage* space and parcellated into 360 regions defined by atlas HCP-MMP1. The mean thickness of each region was as the input of the predictive model. **b**. In one resampling, split all subject into train and test dataset randomly, and estimate the hyperparameters of model in train dataset by grid search with 10-fold cross-validation. Then, retain the model by the tuned hyperparameter, and validate the performance of model by test dataset. To reduce the sampling bias, we repeated 200 times random resampling.

### Prediction model of the memory performance

We designed prediction models for cerebral thickness based on a gradient boosting technique-extreme gradient boosting (XGBoost). XGBoost is an implementation of a gradient boosted trees algorithm that uses a regularized gradient-boosting algorithm to accurately predict the target variable (T. Chen & Guestrin, 2016; Marzi et al., 2023). The model maps the input data to one of the tree’s leaves which contains the score of memory performance that we want to predict. In this study, we implement it using the xgboost package (version 1.6.0, available at https://xgboost.ai) in R (version 4.2.2, available at https://www.r-project.org).

#### MRI features

The mean thickness of 360 regions (corresponding to 360 morphometric features) extracted from the processed T_1_-weighted images are used as the input for the prediction models, and the score of memory performance was used as the predictive target. First, each feature and target are employed as the dependent variable in a linear regression model to estimate the effects of age and education. Subsequently, the residual of each feature and target are standardized using maximum and minimum values of the feature across all subjects, rescaling them into a range from zero to one.

#### Model training and validation

It is commonly necessary to consider preventing overfitting and ensuring a good assessment of the model when it is trained and validated. As illustrated in Figure 1b, we used a nested resampling cross-validation scheme with an outer loop, Monte Carlo cross-validation (MCCV), to reduce variance, and an inner loop, grid search with 10-fold cross-validation, to identify the most optimized hyperparameters. In detail, we fisrt defined a series of hyperparameters first based on previous experiments. Then, we estimated model performance for each hyperparameter using 10-fold cross-validation in the inner loop. The tuned hyperparameter and the entire training set were used to train the optimized model, which was then tested on the test set. In the outer loop, we shuffled the entire dataset and divided it into a training set (80% of the whole set) and a test set (the remaining 20%). Finally, this procedure was repeated 200 times.

#### Model evaluation

To validate the model’s validity, we constructed a mean model as the benchmark in each resampling procedure. We then obtained 200 performance metrics (mean squared error, MSE) for both the prediction model and the corresponding benchmark model.

#### Model interpretation

For the model interpretation, we calculated the feature contribution using the feature importance of XGBoost, which measures feature contributions for the improvement in model performance as the information gain in the model optimization process (Marzi et al., 2023). We selected the top 10% features (regions) as the key regions of the predictive model, and analyzed these key regions as follows. Meanwhile, we overlapped the key regions with a functional network parcellation (Thomas Yeo et al., 2011) and conducted meta-analytical decoding using the *Neurosynth* database (https://www.neurosynth.org) to explore their location and cognitive progression/functional characterization.

### Statistical analysis

A two-sample t-test was used to estimate the sex difference of memory performance and cerebral thickness. Further, a strict multiple comparison correction (Bonferroni) was used for sex differences in cortical thickess. A paired-sample t-test was used to compare the predictive model with the benchmark and to estimate the effectiveness of training. The Pearson correlation was used to estimate the relationship between the thickness of key regions and memory performance. The effect of age and education were controled for in thickness and memory examination scores. The level of statistically significant was set at *p* < 0.05 in all statistical comparison.

## Results

### Demographic information and sex difference in verbal/visuospatial memory performance and cerebral thickness

The demographic characteristics of the participants in this study are presented in Table 1. The study included a total of 1153 participants, with an average age of 66.58 years (65.81 years for females and 67.95 years for males). The study comprised 738 females (64.01%) and 415 males (35.99%), with average education levels of 11.4 years (11.2 years for females and 11.98 years for males). The female group had lower age and education levels compared to the male group (age: *t* = -4.735, *p* < 0.001; education: *t* = -4.017, *p* < 0.001). Therefore, age and education were controlled for in subsequent analyses. There was no significant difference in general cognition performance between females and males (*t* = 1.783, *p* = 0.075).

As shown in Table 1, females performed better than males in verbal memory (*t* = 5.655, *p* < 0.001), while males deomstrated superior performance in visuospatial memory compared to females (*t* = -3.015, *p* = 0.003). These sex differences remained significant even after controlling for the effects of age and education (Figure 2a; verbal memory: *t* = 5.431, *p* < 0.001; visuospatial memory: *t* = -3.201, *p* = 0.001).

**Figure 2.**
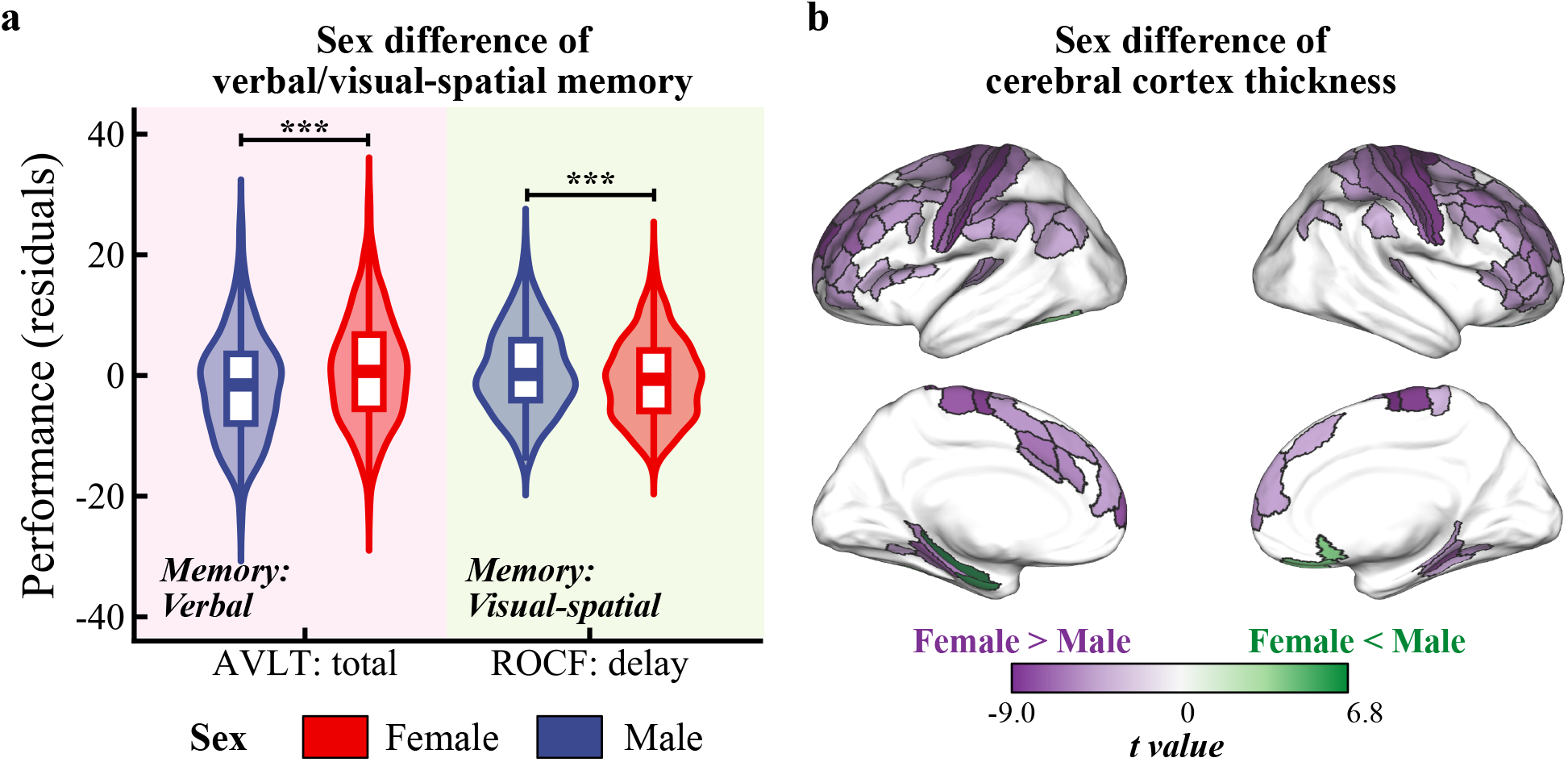
Sex difference of two kind of memory performance (a) and cerebral thickness (b). *Notes*: AVLT, auditory verbal learning test; ROCF, Rey–Osterrieth complex figure test.

There are widespread differences in cortical thickness between males and females, mainly manifested as females having significantly higher cerebral thickness in more regions than males (Figure 2b). Females had a thicker cortex than males in a wide range of frontal, parietal, and some regions of the medial temporal lobe, while males were thicker than females only in the right medial orbitofrontal, left parahippocampal, and left lingual gyrus regions.

### Cerebral thickness can predict verbal and visuospatial memory

First, we evaluated the model performance in predicting verbal memory. Both the mean squared error (MSE) of the predictive model for females and males were significantly lower than those of the benchmark model (female: *t* = -16.343, *p* < 0.001; male: *t* = -10.629, *p* < 0.001). These results indicate that XGBoost, utilizing cerebral cortical thickness as a predictor, effectively predicts verbal memory performance. In addition, we also used DK (Desikan et al., 2006) and BNA (Fan et al., 2016) parcellation atlas to extract cortical thickness and train the model. The results are also consistent (Supp. Table 1).

Next, we calculated the gain of the model and identified the key regions as the top 10% of regions in the gain distribution. The threshold for females was 0.0049 (Figure 3b), while it was 0.0076 for males (Figure 3f).

**Figure 3.**
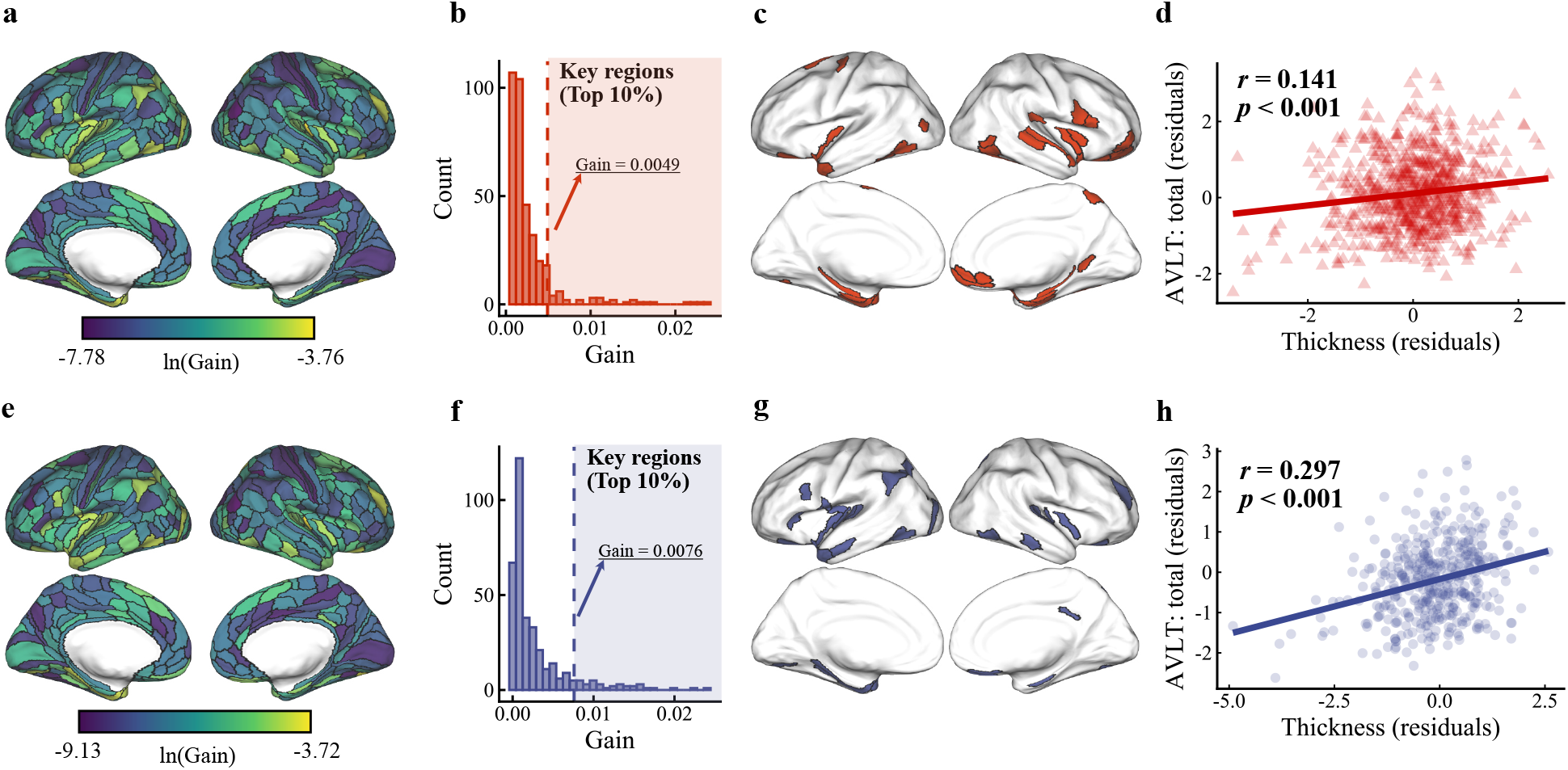
The details about verbal memory predictive model. Top row shown the details information of the model trained in female. From left to right, they are gain value derived from the model, the distribution of gain value across all regions, the key regions defined top 10% regions in the distribution of gain, and the relationship between thickness of key regions and the verbal memory performance. Bottom row shown the same details information of the model trained in male.

Subsequently, we examined the association between the thickness of the key regions and verbal memory performance while controlling for age and education. We found a significant positive relationship between the thickness of the key regions and verbal memory performance (*r* = 0.141, *p* < 0.001 in females, Figure 3d; *r* = 0.297, *p* < 0.001 in males, Figure 3h).

Similarly, for visuospatial memory, we observed that the MSE of the predictive model was significantly lower than that of the benchmark model for both females and males (female: *t* = - 6.608, *p* < 0.001; male: *t* = -7.641, *p* < 0.001). For females, the threshold value for key regions was 0.0064 (Figure 4b), while it was 0.0066 for males (Figure 4f). The thickness of the key regions exhibited a significant positive relationship with visuospatial memory performance in both females and males (*r* = 0.089, *p* = 0.016 in females, Figure 4d; *r* = 0.211, *p* < 0.001 in males, Figure 4h).

**Figure 4.**
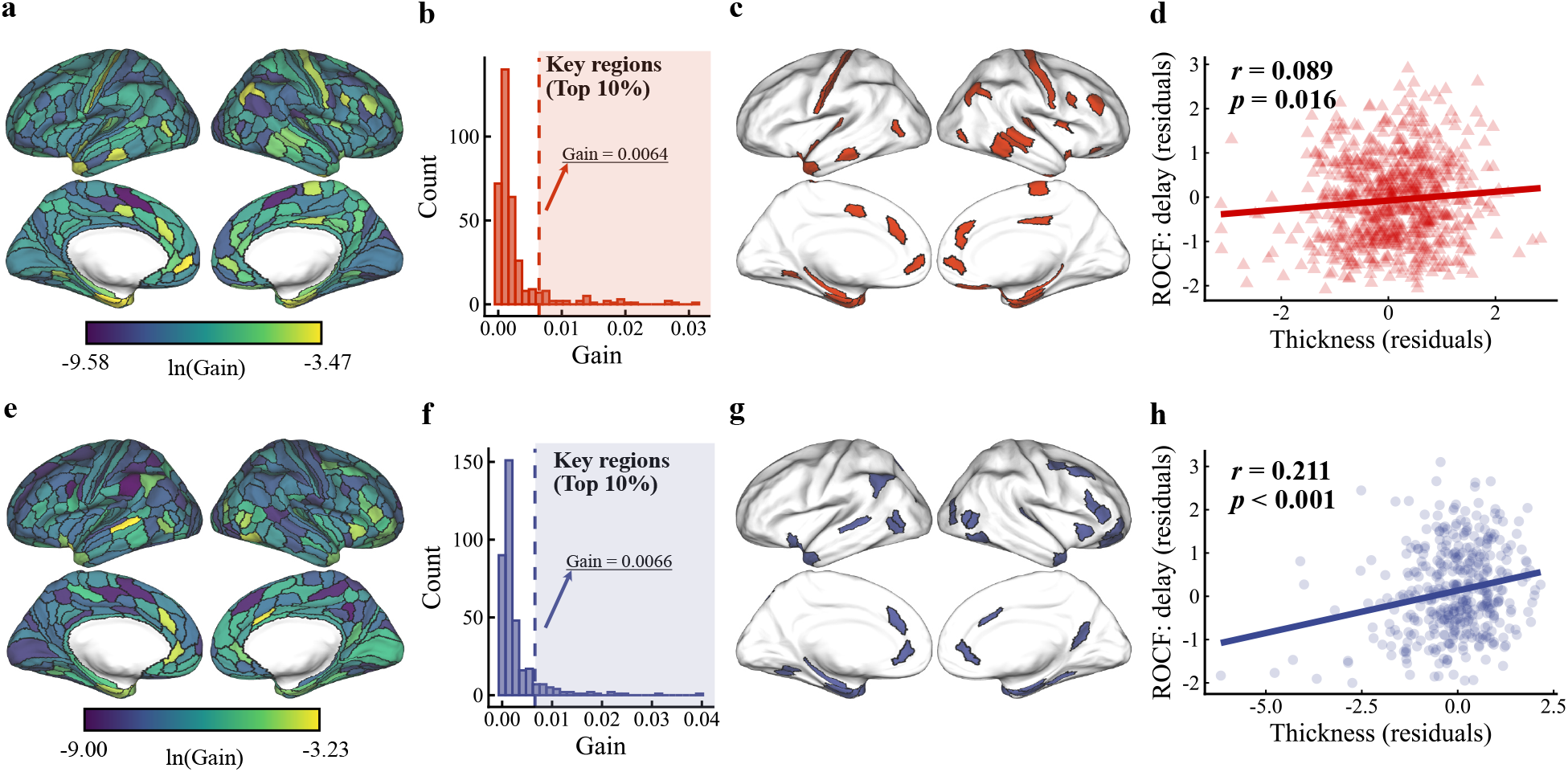
The details about visuospatial memory predictive model. Top row shown the details information of the model trained in female. From left to right, they are gain value derived from the model, the distribution of gain value across all regions, the key regions defined top 10% regions in the distribution of gain, and the relationship between thickness of key regions and the visuospatial memory performance. Bottom row shown the same details information of the model trained in male.

### Sex difference of regions related verbal memory

Based on the predictive model and its feature importance, we defined key regions by selecting the top 10% regions in the feature importance distribution for females and males (Figure 3c and Figure 3g) and categorized them into three groups: female-specific (exclusive to females), male-specific (exclusive to males), and overlapped regions (common regions shared by both sexes). We then overlapped these key regions with Yeo’s 7 network parcellation and calculated the overlap rate for each functional network (Figure 5b). All key regions are publicly available (https://github.com/sangfengchn/MemorySexDifference/tree/main).

**Figure 5.**
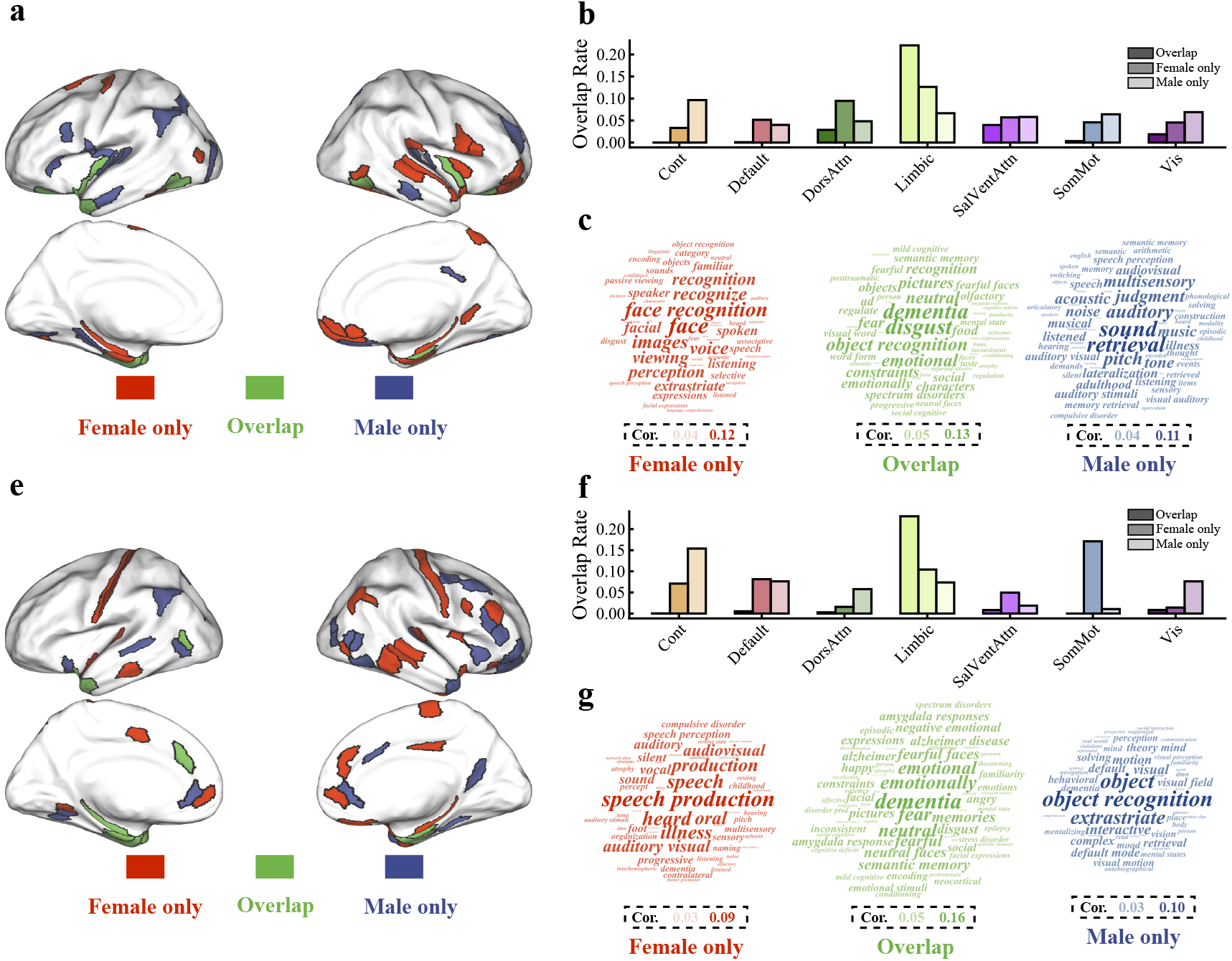
Sex differences in the spatial distribution of key brain regions and its’ decoding function. Top row shown the key regions associated with verbal memory (a, b and c), and bottom row shown the key regions associated with visuospatial memory (e, f and g).

We utilized *Neurosynth*, a well-validated and publicly available platform, for cognitive annotation of these key regions (Figure 5a). Specifically, we input the female-specific, male-specific, and overlapped key regions into *Neurosynth* datebase to decode the related cognitive annotations (“terms”).

The distribution of key brain regions associated with verbal memory is depicted in Figure 5c. We subsequently categorized these brain regions into three types: female-specific, male-specific, and overlapped regions. The specific names and types of these brain regions are presented in Table 2. The overlap rates of these three kinds of brain regions with the seven functional network partitions are illustrated in Figure 5b. The results indicate that brain regions common to both sexes and the limbic system exhibited the highest overlap rates. The female-specific brain regions showed higher overlap rates with the dorsal attention network and the default network, while the male-specific key brain regions demonstrated higher overlap rates with the frontoparietal control network, somatomotor network, and visual network.

**Table 2.**
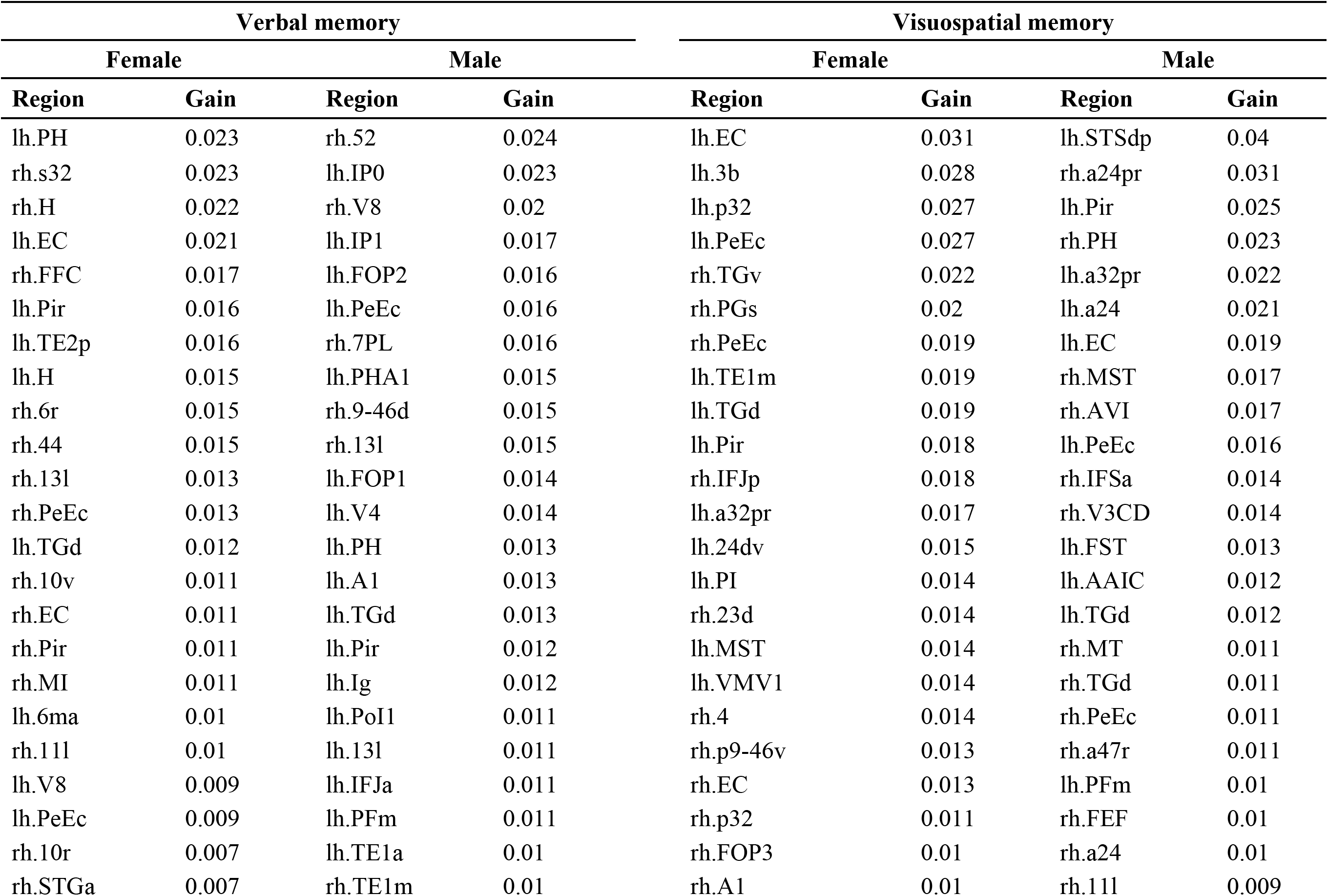
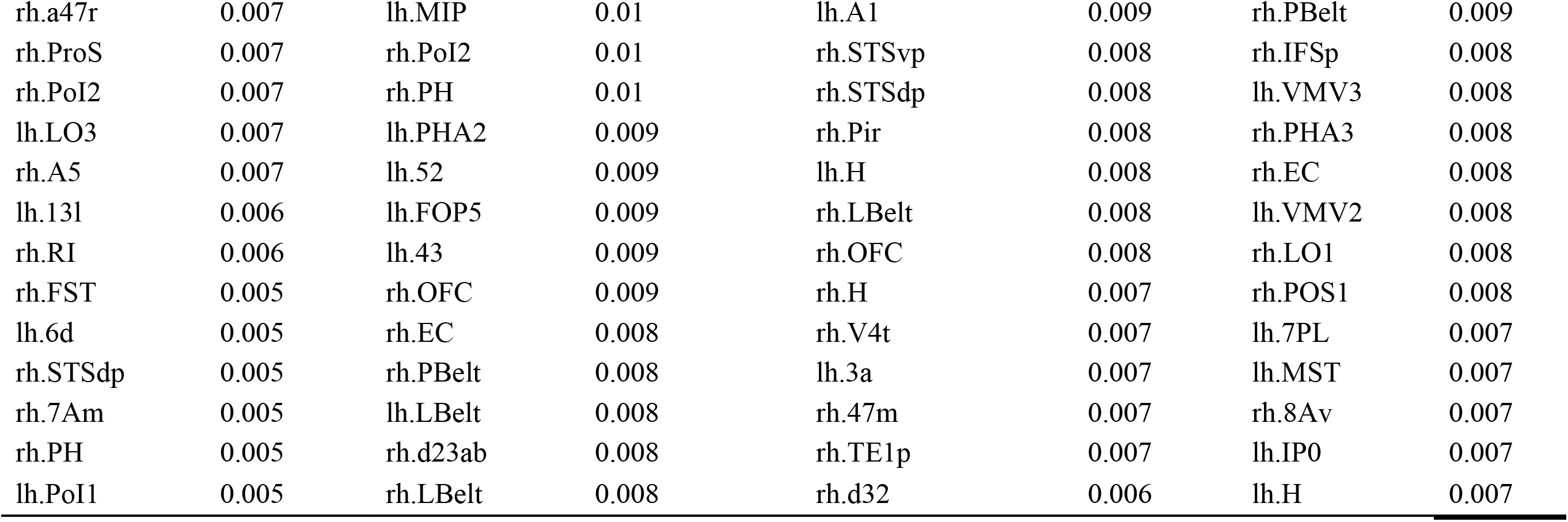
Key regions related verbal memory and visuospatial memory in female and male.

Decoding the key regions using the *Neurosynth* database revealed associations (Figure 5c) between the regions common to both sexes and terms such as “disgust”, “dementia”, and “object recognition”. Female-specific regions were related to terms like “face”, “face recognition”, and “image”, while male-specific regions were associated with terms like “sound”, “retrieval”, and “auditory”. Further detailed information is provided in Table 3. The spatial distribution of key brain regions linked to verbal memory and the associated functional differences between males and females may indicate distinct processing strategies and approaches to verbal memory in each gender.

**Table 3.** Top 20 terms related to key regions of verbal memory and visuospatial memory for female and male.

| Verbal memory |  |  |  | Visuospatial memory |  |  |  |
| --- | --- | --- | --- | --- | --- | --- | --- |
| Female |  | Male |  | Female |  | Male |  |
| analysis | *r* | analysis | *r* | analysis | *r* | analysis | *r* |
| recognition | 0.172 | semantic memory | 0.092 | dementia | 0.164 | dementia | 0.119 |
| faces | 0.169 | objects | 0.090 | progressive | 0.094 | object recognition | 0.115 |
| word form | 0.133 | retrieval | 0.084 | fear | 0.091 | sentence | 0.109 |
| visual word | 0.132 | dementia | 0.084 | emotional | 0.087 | abstract | 0.100 |
| familiar | 0.130 | judgment | 0.082 | neutral | 0.084 | comprehension | 0.095 |
| face recognition | 0.130 | semantic | 0.076 | alzheimer disease | 0.082 | read | 0.092 |
| characters | 0.125 | disgust | 0.073 | atrophy | 0.081 | fear | 0.081 |
| encoding | 0.122 | constraints | 0.067 | constraints | 0.080 | object | 0.081 |
| facial | 0.121 | ad | 0.065 | alzheimer | 0.078 | emotional | 0.079 |
| neutral | 0.118 | encoding | 0.064 | fearful | 0.074 | sentence comprehension | 0.079 |
| images | 0.116 | construction | 0.063 | disgust | 0.073 | neutral | 0.075 |
| recognize | 0.115 | visual | 0.060 | aphasia | 0.073 | mood | 0.073 |
| disgust | 0.113 | sound | 0.059 | emotionally | 0.073 | syntactic | 0.072 |
| objects | 0.111 | matching | 0.058 | amygdala responses | 0.069 | semantic | 0.071 |
| viewing | 0.108 | memories | 0.057 | emotions | 0.068 | extrastriate | 0.070 |
| fear | 0.107 | olfactory | 0.056 | neutral faces | 0.066 | alzheimer disease | 0.068 |
| emotional | 0.105 | bilinguals | 0.053 | pictures | 0.066 | language comprehension | 0.068 |
| expressions | 0.103 | progressive | 0.053 | semantic memory | 0.065 | emotions | 0.066 |
| pictures | 0.099 | chinese | 0.052 | expressions | 0.064 | alzheimer | 0.066 |
| form | 0.097 | switching | 0.052 | mild cognitive | 0.062 | memories | 0.066 |

### Sex difference of regions related visuospatial memory

Using the same analysis method, we analyzed the key brain regions related to visuospatial memory. The spatial distribution of these key brain regions is shown in Figure 4c and Figure 4g. The specific brain region names and kinds are shown in Table 2, and the overlap rates of the three types of brain regions and the seven networks functional parcellation are shown in Figure 5f. The results show that the highest overlap rates were found in brain regions common to both sexes and in the limbic system, which is consistent with key brain regions related to verbal memory. However, female-specific brain regions were more often located in the somatosensory-motor and ventral attention networks, whereas male-specific key brain regions had higher overlap rates with the dorsal attention network, frontoparietal control network and visual network. Similarly, decoding using the *Neurosynth* database revealed (Figure 5g and Table 3) that regions common to both sexes were associated with “dementia”, “fear”, and “emotionally”, female-specific regions were associated with “speech production”, “speech”, and “heard”, and male-specific regions were associated with “object recognition”, “object”, and “interactive”. The spatial distribution of key brain regions associated with visuospatial memory and associated functional differences between males and females suggest that males and females may be processing such information with different brain pathways and processing strategies.

## Discussion

In this study, we employed an interpretable machine learning model, utilizing nested random sampling and grid search techniques, to train predictive models separately for females and males. These models were used to analyze the relationship between brain structural measures and the performances of two types of memory in a sample of 1153 older adults. The study identified the key brain regions associated with verbal and visuospatial memory in each sex, based on the predictive models. The main results of this study are as follows: (1) In the elderly non-dementia population, females outperformed males in verbal memory, while males outperformed females in visuospatial memory. (2) Females had thicker cortex than males in a wide range of frontal, parietal, and some regions of the medial temporal lobe. In constrast, males were thicker than females only in the right medial orbitofrontal, left parahippocampal, and left lingual gyrus regions. (3) The cortical thickness in both males and females could accurately predict verbal memory and visuospatial memory scores. Furthermore, the cortical thickness of key brain regions predicted by the model was significantly correlated with the respective memory scores. (4) Overlapping and decoding analysis revealed that the key brain regions associated with female-superior verbal memory mainly located in limbic, dorsal attention and default mode networks. These regions are also associated with face recognition and perception processes. Meanwhile, the key brain regions associated with male-superior visuospatial memory are mainly located in dorsal attention, frontoparietal and visual networks, and associated with object recognition processes.

The present study verified the superior performance of females in verbal memory and males in visuospatial memory in an elderly population. Sex differences in cortical thickness were also verified. Specifically, females had significantly thicker cortices than males in most areas of the superior part of the brain (including the frontal and parietal lobes) and parts of the medial temporal lobe. In contrast, males had thicker cortices than females in the right medial orbitofrontal area and the left parahippocampal cortex. These results were generally consistent with previous studies (Ritchie et al., 2018; Sowell et al., 2007). However, sex differences in cortical thickness did not correspond to sex differences between the two types of memory performance: not all brain regions with differences were associated with memory performance. We therefore consider that the relationship between brain structure and the two types of memory may be nonlinear, resulting from multiple brain regions acting together, rather than each brain region contributing equally to verbal and visuospatial memory performance. From this perspective, the multi-variables or brain network analysis may be more suitable methods to examine the relationship between brain structure and cognitive performance. Ge et al. (2021) used cortical morphological networks derived from multivariate modeling of regional structural measurs to examine sex difference, and found different pattern of sex differences compared to sex differences in structural measures alone. In this study, we used a multivariate, data-driven method to explore the sex differences in the relationship between brain structural measures and memory performance.

The results of the present study suggest that the key brain areas associated with both female-superior verbal memory and male-superior visuospatial memory are involved in high-level, abstract, and holistic cognitive processes. In contrast, the brain areas associated with female-inferior visuospatial memory and male-inferior verbal memory performance are more involved in low-level, concrete, and specific cognitive processes. This pattern seems to indicate that superior memory performance generally occurs in brain regions associated with abstract cognition. The findings also imply sex differences in the brain extend beyond concrete indicators to differences in brain-behavior relationships. Therefore, when researching brain-behavior links, conducting separate within-sex analyses may be an important consideration for future studies.

Although it is widely accepted that females have an advantage in verbal-related abilities, studies have also reported the different females-superior patterns in different verbal abilities (Toivainen et al., 2017). In a meta-analysis (Hirnstein et al., 2022), females showed superior performance in phonological fluency rather than semantic fluency, and outperformed males in verbal recall and recollection. However, the female advantage in verbal working memory performance was not significant. The verbal memory performance assessed in the present study was characterized by scores on an auditory verbal learning test (AVLT), which measures verbal recall ability. However, this study focused on exploring sex differences in the relationship between brain structure and verbal memory and was limited in delineating and analyzing different aspects of verbal memory in detail due to the cohort design.

Previous studies have pointed out that differences in spatial abilities between females and males arise not only from biological (especially hormonal) and social factors, but are also related to differences in visual perception processes between females and males (Qian et al., 2022). Males have higher sensitivity to visual stimuli compared to females (Abramov et al., 2012; Brabyn & McGuinness, 1979; Qian et al., 2022). The more sensitive visual perception in males allows for better spatial information processing and thus superior performance in spatial memory tasks. This may explain why males outperform females in examination of visuospatial memory. The present study indeed found differences in brain regions involved in visuospatial memory between females and males in terms of brain structure. This may account for the different visual perception processes between males and femelas. More fundamentally, the basis for these sex differences may be directly linked to gonadal steroid hormones (Abramov et al., 2012).

In addition to cortical thickness, we also performed the same analysis on other measures of the cerebral cortex, including cortical surface area and volume (Supp. Table 1). We found that cortical surface area did not predict verbal memory scores in the female population, while cortical volume results were consistent with those for cortical thickness and significantly predicted corresponding memory performance in all four groups (sex * two perfromances). However, considering that cortical volume is a mixed index obtained from cortical thickness and surface area (Panizzon et al., 2009), the present study finally analyzed and reported using cortical thickness as the main study measure. The radial unit hypothesis of cortical development postulates that the size of the cortical surface area is driven by the number of cells ontogenetic columns, whereas cortical thickness is influenced by the number of cells within a columns (Rakic, 1988). Our findings indicated that cortical thickness is a more sensitive predictor of verbal and visuospatial memory performance in both females and males.

## Limitations

While we conducted extensive analyses and discussion, there are inherent limitations to the present study that must be acknowledged. First, as an exploratory study examining the relationship between brain structure and memory performance, the present study can only speculate about possible associations and does not determine causality regarding how brain structure impacts verbal or visuospatial memory in females and males. Second, since the data included in this study were all from the BABRI study cohort, the subjects in this study were all from the elderly population. Although we controlled for age in our analyses to mitigate the effect of age as much as possible, the current findings may still have limited generalizability to other age groups. Finally, given the sample size and computational costs, only 200 random samples were conducted in this study. Although more than a thousand subjects were included in total in this study, after splitting into sex groups, only around half of the subjects were in each group. In the future, more robust results could be obtained by combining multiple large imaging databases. Of course, recruiting more subjects would require more random sampling, so computational resources would be an issue to consider.

## Conclusion

This study is based on a large sample of community-dwelling elderly participants and utilizes a machine learning model to explore sex differences in verbal/visuospatial memory as well as associated sex differences in related cognitive processes and spatial distribution of relevant brain regions. The results confirm the superior performance of females in verbal memory and males in visuospatial memory. Additionally, differences are found in the brain regions associated with memory performance between females and males. Using *Neurosynth* decoding and overlapping with Yeo’s 7 networks, we further discovered that the brain regions related to sex-specific superior memory performance of females and males are mainly located in the dorsal attentional network and involved in more advanced and holistic cognitive processes are in both females and males. In addition, the key regions are also located in the default mode network in females and the visual network in males.

## Authors’ Contributions

F.S. contributed to the research ideas, data analysis, results visualization, and drafting and final approval of the manuscript. S.Z. contributed to the data analysis, results interpretation, and manuscript revision. Z.L. contributed to the results visualization, results interpretation, and manuscript revision. Y.Y. contributed to the results interpretation and manuscript revision. Y.C. contributed to the results interpretation, manscript revision, and final approval of the manuscript. She is also the guarantor of this work. Z.Z. is the primary leader of the BABRI cohort and is the lead applicant for the funding required for this study.

## Acknowledgement

This work was supported by International Cooperation and Exchange of the National Natural Science Foundation of China (grant number 81820108034, https://www.nsfc.gov.cn/english/site_1/policy/B2/2017/12-29/59.html), National Natural Science Foundation of China (grant number 82071205, and 8213000253, https://www.nsfc.gov.cn), and State Key Program of National Natural Science of China (grant number 81430100, https://www.nsfc.gov.cn/Portals/0/fj/english/fj/pdf/2013/021.pdf). The BABRI team put a lot of effort into subject recruitment, neuropsychological testing, and data organization. All subjects of this research also contributed.

