## Supplementary material for "Cerebral thickness reveals sex differences in verbal and visuospatial memory": Supp. Table 1

### Supplementary materials

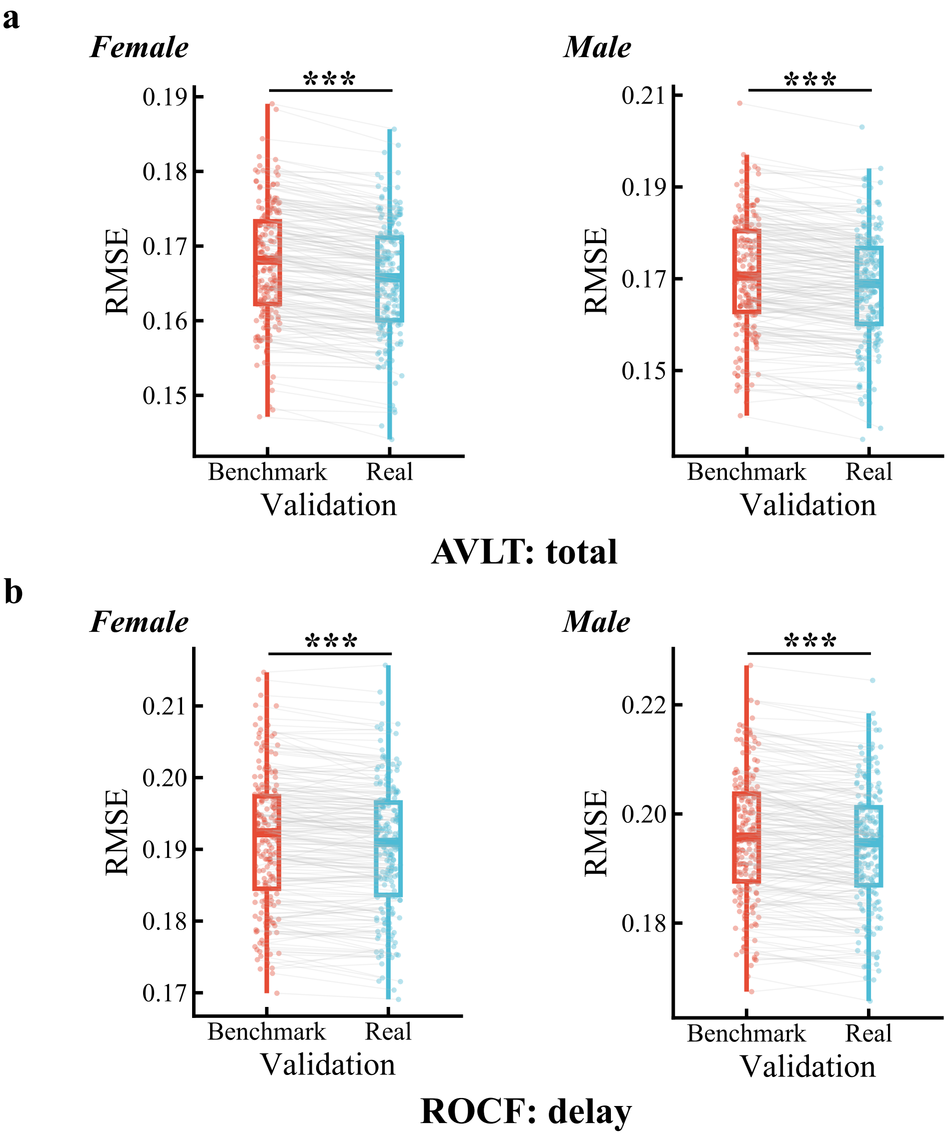


Supp. Figure 1. The performance of the predicitive models trained on cortical thickness using HCP MMP 1.0 parcellation and benchmark models on the test set.

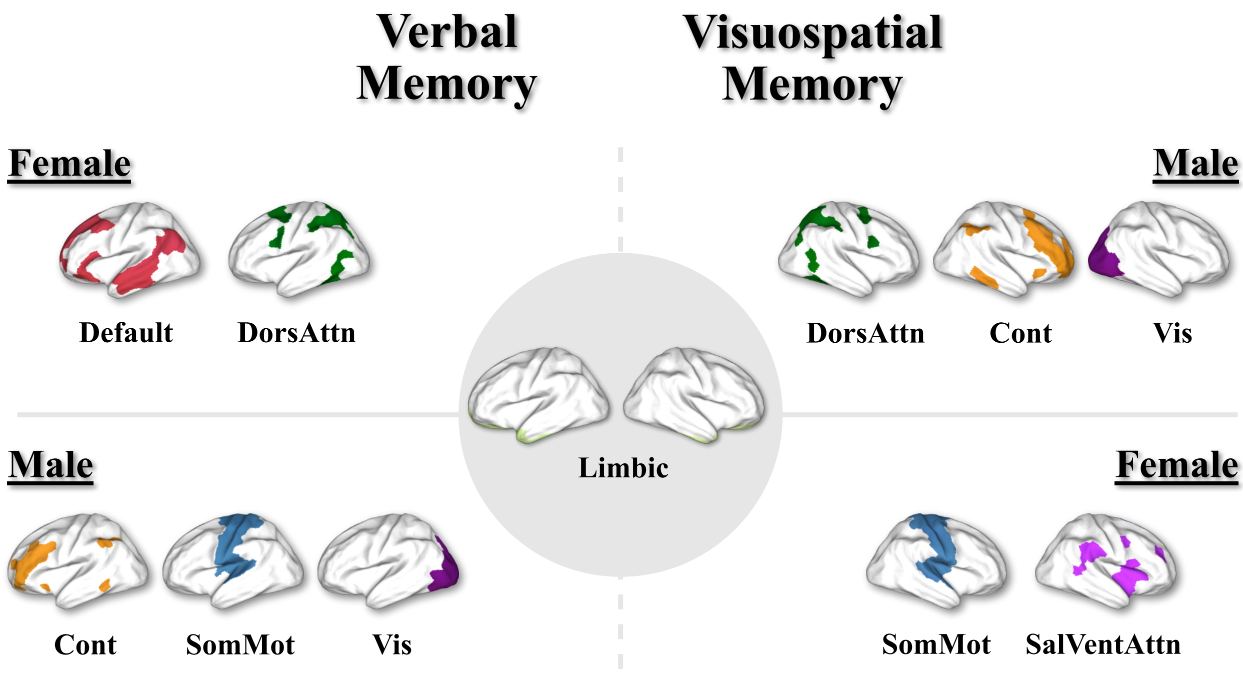


Supp. Figure 2. Sex differences in the spatial distribution of key brain regions associated with verbal and visuospatial memory performance.

Supp. Table 1. The performance of the predicitive models trained on cortical measures using HCP MMP 1.0 parcellation and benchmark models on the test set.

| Measures | Cognition/Sex | Root Mean Square Error (RMSE) | | *t(df)* | *p* | *95% CI* |
|---|---|---|---|---|---|---|
| | | Real (M±SD) | Benchmark (M±SD) | | | |
| **Thickness** | ***AVLT: total*** | | | | | |
| | Female | 0.1657±0.0075 | 0.1678±0.0078 | -16.343(199) | <0.001 | [-Inf, -0.0019] |
| | Male | 0.1684±0.0119 | 0.1708±0.0122 | -10.629(199) | <0.001 | [-Inf, -0.0021] |
| | ***ROCF: delay*** | | | | | |
| | Female | 0.1904±0.0090 | 0.1912±0.0092 | -6.608(199) | <0.001 | [-Inf, -0.0006] |
| | Male | 0.1940±0.0111 | 0.1958±0.0116 | -7.641(199) | <0.001 | [-Inf, -0.0014] |
| **Area** | ***AVLT: total*** | | | | | |
| | Female | 0.1675±0.0075 | 0.1678±0.0078 | -2.436(199) | 0.008 | [-Inf, -0.0001] |
| | Male | 0.1693±0.0119 | 0.1708±0.0122 | -9.199(199) | <0.001 | [-Inf, -0.0012] |
| | ***ROCF: delay*** | | | | | |
| | Female | 0.1905±0.0089 | 0.1912±0.0092 | -5.679(199) | <0.001 | [-Inf, -0.0005] |
| | Male | 0.1965±0.0112 | 0.1958±0.0116 | 4.557(199) | >0.999 | [-Inf, 0.0011] |
| **Volume** | ***AVLT: total*** | | | | | |
| | Female | 0.1663±0.0074 | 0.1678±0.0078 | -13.409(199) | <0.001 | [-Inf, -0.0013] |
| | Male | 0.1660±0.0121 | 0.1708±0.0122 | -22.467(199) | <0.001 | [-Inf, -0.0045] |
| | ***ROCF: delay*** | | | | | |

| | | | | | |
|---|---|---|---|---|---|
| Female | 0.1899±0.0089 | 0.1912±0.0092 | -10.257(199) | <0.001 | [-Inf, -0.0011] |
| Male | 0.1948±0.0112 | 0.1958±0.0116 | -5.219(199) | <0.001 | [-Inf, -0.0007] |

Supp. Table 2. The performance of the predicitive models trained on cortical thickness using DK and BNA parcellation and benchmark models on the test set.

| Atlas | Cognition/Sex | Root Mean Square Error (RMSE) | | *t(df)* | *p* | *95% CI* |
|---|---|---|---|---|---|---|
| | | **Real (M±SD)** | **Benchmark (M±SD)** | | | |
| **DK** | ***AVLT: total*** | | | | | |
| | Female | 0.1651±0.0077 | 0.1668±0.0078 | -12.689(199) | <0.001 | [-Inf, -0.0015] |
| | Male | 0.1650±0.0124 | 0.1689±0.0132 | -17.018(199) | <0.001 | [-Inf, -0.0036] |
| | ***ROCF: delay*** | | | | | |
| | Female | 0.1914±0.0086 | 0.1918±0.0086 | -2.448(199) | 0.015 | [-Inf, -0.0001] |
| | Male | 0.1899±0.0123 | 0.1925±0.0125 | -11.700(199) | <0.001 | [-Inf, -0.0022] |
| **BNA** | ***AVLT: total*** | | | | | |
| | Female | 0.1650±0.0074 | 0.1678±0.0078 | -20.682(199) | <0.001 | [-Inf, -0.0025] |
| | Male | 0.1679±0.0119 | 0.1708±0.0122 | -12.734(199) | <0.001 | [-Inf, -0.0025] |
| | ***ROCF: delay*** | | | | | |
| | Female | 0.1910±0.0089 | 0.1912±0.0092 | -1.838(199) | 0.034 | [-Inf, <-0.0001] |
| | Male | 0.1924±0.0109 | 0.1958±0.0116 | -14.015(199) | <0.001 | [-Inf, -0.0029] |

Note. DK, Desikan-Killiany Atlas; BNA, Brainnetome Atlas.
